# Inhibition of the amino-acid transporter LAT1 demonstrates anti-neoplastic activity in medulloblastoma

**DOI:** 10.1101/428722

**Authors:** Yann Cormerais, Marina Pagnuzzi-Boncompagni, Sandra Schrötter, Sandy Giuliano, Eric Tambutté, Hitoshi Endou, Michael F. Wempe, Gilles Pagès, Jacques Pouysségur, Vincent Picco

**Affiliations:** Centre Scientifique de Monaco, Biomedical Department, 8 Quai Antoine Ier, MC-98000 Monaco, Principality of Monaco; Department of Genetics and Complex Diseases, Harvard T. H. Chan School of Public Health, Boston, MA, USA; Centre Scientifique de Monaco, Marine Biology Department, 8 Quai Antoine Ier, MC-98000 Monaco, Principality of Monaco; J-Pharma, Co. Ltd., Yokohama, Japan; School of Pharmacy, Anschutz Medical Campus, University of Colorado Denver, Aurora, Colorado 80045; University of Cote d’Azur, Institute for Research on Cancer and Aging of Nice (IRCAN), CNRS UMR 7284; INSERM U1081, Centre Antoine Lacassagne, France

**Keywords:** Medulloblastoma, Amino acid transport, GCN2, ATF4, mTORC1

## Abstract

Most cases of medulloblastoma (MB) occur in young children. While the overall survival rate can be relatively high, current treatments combining surgery, chemo- and radiotherapy are very destructive for patient development and quality of life. Moreover, aggressive forms and recurrences of MB cannot be controlled by classical therapies. Therefore, new therapeutic approaches yielding good efficacy and low toxicity for healthy tissues are required to improve patient outcome. Cancer cells sustain their proliferation by optimizing their nutrient uptake capacities. The L-type amino acid transporter 1 (LAT1) is an essential amino acid carrier overexpressed in aggressive human cancers that was described as a potential therapeutic target. In this study, we investigated the therapeutic potential of JPH203, a LAT1-specific pharmacological inhibitor, on two independent MB cell lines belonging to subgroups 3 (HD-MB03) and Shh (DAOY). We show that while displaying low toxicity towards normal cerebral cells, JPH203 disrupts AA homeostasis, mTORC1 activity, proliferation and survival in MB cells. Moreover, we demonstrate that a long-term treatment with JPH203 does not lead to resistance in MB cells. Therefore, the present study suggests that targeting LAT1 with JPH203 is a promising therapeutic approach for MB treatment.

## INTRODUCTION

Medulloblastoma (MB) is the most prevalent pediatric brain tumor [1]. Recent advances in genetic characterization of the disease have led to an international consensus classification of MB into four biologically and clinically relevant subtypes: the wingless (Wnt), the sonic hedgehog (Shh), and the more similar though molecularly distinguishable groups 3 and 4 [2,3]. The most recent studies combining genetics, epigenetics and RNA expression data have refined and further delineated several sub-groups within these groups [3–6]. MB subgrouping is used to orient the therapeutic approach that comprises surgical removal of the tumor, cranio-spinal radiation therapy (RT) and chemotherapy. Shh and Wnt subgroups show the highest overall survival while group 4 and, to a higher extent, group 3 MBs are of poor prognosis with a strong tendency to form metastasis [2,3]. Although the overall survival at 5 years reaches 70% in all MB types taken together, even peaking over 90% in the Wnt group, toxic effects due to the treatments often lead to irreversible damages that severely hamper the children’s cognitive development and general quality of life. In particular, RT dose and age of exposure to RT are well-documented risk factors for long-term cognitive impairments in MB survivors [7–9]. Reducing RT doses has been proposed however it is unsuitable for high risk MBs [8,9]. Moreover, recurrences are fatal in a large majority of cases and are characterized by strong divergence of the relapsed tumors as compared to the initial tumors [10]. A challenge therefore lies in finding new treatments that combine high efficacy and little toxicity. Personalized targeted therapy appears to be a promising approach in this context [11,12]. Clinical trials involving Shh pathway targeted inhibitors have enrolled Shh group MB patients based on this assertion. To date, these trials have not led to the expected results as they have induced unbearable toxicity and tumor resistance to the treatment [13–15]. All these features support the need to develop and design novel therapies against alternative targets.

Rapidly growing tumors experience an increased demand for nutrients in order to sustain their proliferative metabolism. Up-regulation of processes related to the supply of glucose, amino acids and lipids is thus a hallmark of cancer metabolism [16,17]. Essential amino acids (EAAs), which cannot be synthetized *de novo* by human cells, are absolutely required for cancer cell proliferation. Indeed, some of these nutrients can also be converted to essential metabolite intermediates of the TCA cycle, further participating in tumor energy metabolism and macromolecule synthesis. Notably, catabolism of the branched chain EAA subclass (leucine, isoleucine, and valine) is required for growth of some brain tumors [18]. In addition, leucine is an essential signaling molecule required to sustain the activation of mTORC1, the master kinase of protein, lipid, nucleotide syntheses and cell proliferation [19,20]. Therefore, considering the high expression of LAT1 in MB cells [21], we hypothesized that targeting EAA uptake might be an innovative strategy for MB treatment. The L-type amino acid transporter 1, LAT1 (SLC7A5), is a 12-transmembrane protein responsible for Na+-independent transport of large neutral EAA (Leu, Val, Ile, Phe, Trp, His, Met, Tyr) in tight association with chaperone CD98 (SLC3A2). LAT1 is an obligatory exchanger with the uptake of one amino acid (AA) being coupled to the efflux of another. LAT1 is overexpressed in aggressive human cancers including MB and has been described as a potential therapeutic target [21,22]. In a previous study, we demonstrated that LAT1 activity is essential for tumor growth [23]. Indeed, genetic disruption of LAT1 in colorectal and lung adenocarcinoma cell lines leads to EAA starvation, mTORC1 inactivation and growth arrest [23,24]. In addition, our lab with others demonstrated that treatment with JPH203, a specific inhibitor that targets LAT1 and no other LAT, recapitulates these effects in several cancer cell lines, including glioma cell lines [23,25,26]. However, the efficiency of this compound had never been addressed in MB. In the present study, we demonstrate that LAT1 is the main leucine transporter in two independent cell lines isolated from Shh (DAOY) and Group 3 (HD-MB03) MBs [27,28]. We subsequently show that JPH203 treatment disrupts AA homeostasis, mTORC1 activity as well as proliferation and survival of both MB cell lines. Further, in contrast to its effect on MB cell lines, JPH203 displays low toxicity on two normal cerebral cell types. Finally, we show that although MB cells try to adapt to a JPH203 chronic treatment by upregulating the expression of AA transporters, no resistant clones could be isolated from the cell populations. Altogether our *in vitro* results strongly suggest that JPH203 is a promising therapeutic candidate for treating MB.

## MATERIAL AND METHODS

### Cell culture and pharmacological inhibitor

The murine cerebellar astrocyte cells C8-D1A were obtained from the American Tissue Culture Collection (ATCC^®^ number CRL-2541, passage 6) and cultivated in Dulbecco’s modified Eagle’s medium (DMEM) supplemented with 10% inactivated fetal calf serum (iFCS, Dutscher). The human MB cell lines DAOY (ATCC^®^ number HTB-186, passage 15) and HD-MB03 [27] (Creative Bioarray Cat.No.CSC-C6241X, passage 60) were obtained from Dr. C. Pouponnot (Institut Curie, France) and cultivated in Minimum Essential Medium (MEM, Invitrogen) supplemented with 10% iFCS and 90% RPMI-1640 supplemented with 10% iFCS respectively. Primary murine neurons were isolated from E16.5 mouse embryos cortices dissociated with trypsin as previously described [29]. Briefly, neurons were plated onto 12-wells plates and cultured in neurobasal medium (21103049 Gibco) supplemented with B27 (A3582801 Gibco) and glutamine. All the cells were maintained at 37°C and 5% CO2. The LAT1-targetting pharmacological inhibitor JPH203 (J-Pharma, Yokohama, Japan) was resuspended in dimethyl sulfoxide (DMSO). For long-term chronic treatment with JPH203, DAOY and HD-MB03 cell populations were cultivated in 100mm diameter dishes in Ham’s F12 medium (21765-029 Gibco) supplemented with 20μM and 30μM of JPH203 or DMSO (same amount as the 30μM JPH203 condition). Cells were passaged regularly based on their confluence level and the medium was changed at least every 3 days to avoid degradation of the compound. After trypsinization, cells were seeded at concentrations of 0. 5.10^6^ and 3.10^6^ cells per dish for DAOY and HB-MB03 respectively.

### Immunoblotting

Cells were lysed in 1.5× Laemmli buffer, and protein concentrations were determined using the Pierce BCA protein assay (23227 Thermo Scientific). Protein extracts (40 μg) were separated by electrophoresis on 10% SDS polyacrylamide gel and transferred onto polyvinylidene difluoride membranes (Millipore). Membranes were blocked in 5% nonfat milk in TN buffer (50 mmol/L Tris-HCl pH7.4, 150 mmol/L NaCl) and incubated with the following anti-human antibodies: LAT1 (1:1,000, KE026 TransGenic Inc.), CD98 (1:1,000, SC-9160 Santa Cruz Biotechnology), GCN2 (1:250, sc-374609 Santa Cruz Biotechnology), phospho-GCN2 (1:500, ab75836 Abcam), EIF2α (1:1,000, ab5369 Abcam), phospho-EIF2α (1:1,000, ab32157 Abcam), ATF4 (1:1,000, 11815S CST), phospho-p70-S6K (1:1,000, 9202S CST), p70-S6K (1:1000, 9201S, CST), RPS6 (1:1,000, 2217S CST), phospho-RPS6 (1:1,000, 2215S CST), ERK1/2 (1:10,000 MA5-16308, Thermo Scientific), and tubulin (1:10,000 MA5-16308, Thermo Scientific). Immunoreactive bands were detected with horseradish peroxidase-coupled anti-mouse or anti-rabbit antibodies (Promega) using the ECL system (Merck Millipore WBKLS0500). Acquisition of the immunoblot images were performed using LI-COR Odyssey Imaging System. Quantification of the intensity of the western blot bands were performed with ImageJ software (NIH, USA).

### L-[14C]-Leucine uptake

Cells (2.5 × 10^5^) were seeded in triplicates onto 35-mm dishes. 24 hours later, culture media were removed and cells were carefully washed with prewarmed Na+-free Hank’s Balanced Salt Solution (HBSS: 125 mmol/L choline chloride, 4.8 mmol/L KCl, 1.2 mmol/L MgSO4, 1.2 mmol/L KH2PO4, 1.3 mmol/L CaCl2, 5.6 mmol/L glucose, and 25 mmol/L HEPES), preincubated in 1.0 mL of pre-warmed Na+-free HBSS at 37°C for 5 minutes before adding substrates for the uptake experiment. Cells were then incubated at 37°C for 1 minute in 1 mL of Na+-free HBSS containing 1 μmol/L of L-[14C]-leucine (0.03 μCie/mL; PerkinElmer). Subsequently, cells were washed three times with ice-cold Na+-free HBSS containing 1 mmol/L of non-radiolabeled leucine. Cells were then lysed with 1 mL of 0.1 N NaOH and mixed with 3.5 mL of Emulsifier-Safe cocktail (PerkinElmer). Radioactivity was measured using a β-scintillation counter. For the inhibition experiments, the uptake of 1 μmol/L L-[14C]-leucine is examined in the presence of JPH203 (30μM).

### Proliferation and viability assays

The different cell lines (2.5 ×10^4^ cells for 7 days, 5 × 10^4^ cells for 3 days) were seeded onto 6-well plates in triplicates. We measured proliferation by trypsinizing the cells and counting them daily with a Coulter Z1 (Beckman) after 48 hours. The cell proliferation index was calculated as “fold increase” by standardizing each measurement to the cell number obtained 24 hours after seeding (day 0). Viability assays were performed using an ADAM-MC automatic cell counter (Nanoentek) according to the manufacturer’s protocol. Briefly, cells were seeded in 12-well plates, treated for 48 hours with the indicated concentrations of JPH203 (20 and 30μM) or the corresponding amounts of DMSO (0.2 and 0.3% respectively), trypsinized and the samples were analyzed according to the manufacturer’s protocol.

### Three-dimensional growth assay

DAOY and HD-MB03 3D cultures were prepared using ultra-low attachment 96-well plates (Corning). Cells (5000) were seeded in 200μL medium per well. The spheroids were cultured for 8 days and pictures were taken with an AMG Evos microscope 40x objective (Thermo Fisher Scientific Inc). Spheroid areas were measured using ImageJ 1.51j8 software (National Institute of Health) [30].

### Scratch assay

DAOY cells (800000) were seeded in 60mm diameters and grew until they reached confluency (2 days). A wound was then done in the monolayer by scratching it with a 10μL pipette tip (Starlab). The culture medium was then replaced by fresh medium containing DMSO or the indicated concentrations of JPH203. Paired measurements of the wound width were done at 0 and 8 hours at 6 different places for each condition.

### Quantitative real-time PCR

DAOY and HD-MB03 (250 000 and 500 000 cells respectively) were seeded in 6-well plates in triplicate. The next day, the cells were treated with the indicated concentrations of JPH203 or with 0.3% DMSO, a concentration corresponding to the 30μM JPH203 condition. Total RNA were purified after 72 hours incubation with a Nucloespin RNA purification kit (Macherey-Nagel cat. no. 740955) according to the manufacturer’s instructions. Total RNA samples (1 μg) were used to perform a reverse transcription reaction with a Maxima First Strand cDNA Synthesis kit (Thermo Scientific, cat. no. K1671) according to the manufacturer’s instructions. cDNAs were then diluted 5 times and 5μL of each sample was used for RT qPCR assays. Briefly, the experiments were run on a Step One Plus system (Applied Biosystems cat. no. 4676600) with Thermo Scientific’s Taqman kits targeting SLC7A5/LAT1 (cat. no. Hs00185826), SLC7A8/LAT2 (cat. no. 00794796), SLC43A1/LAT3 (cat. no. Hs00992327) and SLC3A2/LAT4 (cat. no. Hs00379955) according to the manufacturers protocols. RPLP0 (cat. no. Hs9999902) was used as the internal control. Three technical replicates were run for each experiment and at least three independent experiments were performed for each experimental condition.

### Statistical analysis

Data are expressed as mean ± SD. Each experiment was performed at least three times. Statistical analysis was done with the unpaired Student t-test. Differences between groups were considered statistically significant when P < 0.05 in a Student’s t-test.

## RESULTS

### LAT1 is the main Na^+^ independent leucine transporter in HD-MB03 and DAOY MB cell lines and is essential for AA homeostasis and mTORC1 activity

We first demonstrated that LAT1 and its chaperone CD98 are expressed in HD-MB03 and DAOY cell lines (Fig. 1A). The multiple bands observed in the CD98 blot are due to posttranslational modifications of the protein. None of them is detectable in protein extract obtained from CD98 knock-out cells [23]. Functional activity of LAT1 was quantified by measuring the Na^+^-independent rate of leucine transport in the presence or absence of JPH203, a specific LAT1 inhibitor (Fig. 1B). JPH203 completely abolished leucine uptake (Fig. 1B), suggesting that LAT1 is the main functional Na_+_ independent leucine transporter in these two MB cell lines. Next, we investigated the effects of LAT1 inhibition on the two AA-sensing pathways: GCN2 and mTORC1 (Fig.1C) [31]. In both cell lines, LAT1 inhibition resulted in the activation of the AA stress response pathway GCN2, observed through increased phosphorylation of GCN2 and EIF2α and upregulation of ATF4 expression (Fig.1C and Fig.S1). Moreover, JPH203 treatment resulted in a strong decrease in mTORC1 activity, scored by the phosphorylation of its two effectors: p70-S6K1 and the ribosomal protein S6 (Fig.1C and Fig.S1). Altogether these results demonstrate that JPH203 treatment leads to AA starvation and suggest that LAT1 activity is required for AA homeostasis in cells belonging to

**Figure 1:**
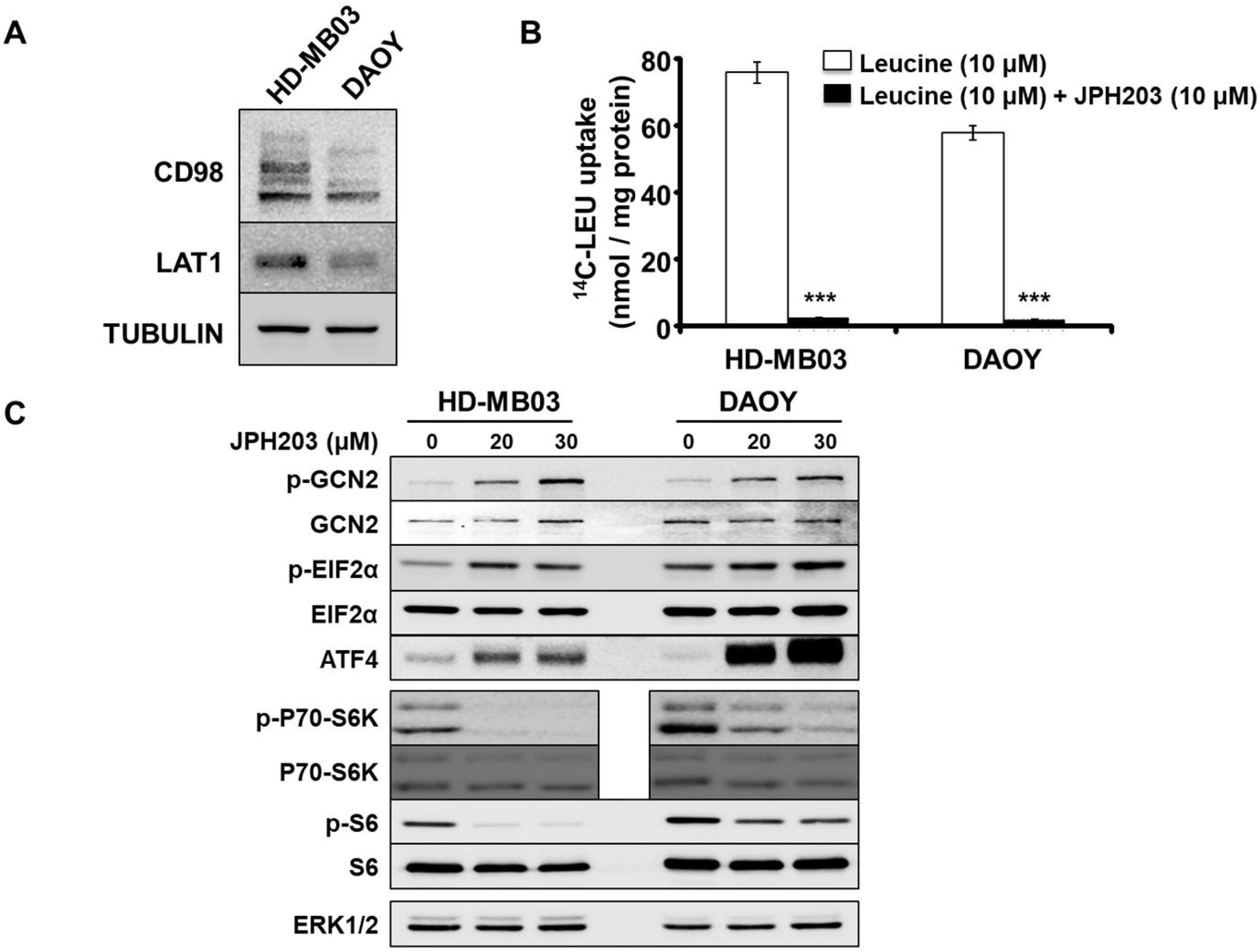
LAT1 is the main leucine Na_+_ independent transporter expressed in MB cell lines HD-MB03 and DAOY and is essential for AA homeostasis and mTORC1 activity. (A) Western blot analysis of the expression levels of LAT1 and its chaperone CD98 in HD-MB03 and DAOY. Tubulin served as a loading control. The results presented are representative of at least three independent experiments (B) Transport assay using radiolabelled leucine (^14^C-LEU) in the absence or presence of 10μM of JPH203 (***: p<0.001, Student’s t-test). (C) Activity of the two AA sensing pathways GCN2 and mTORC1 were analyzed by immunoblot in the absence or presence of either 20 or 30μM of JPH203. ERK1/2 served as a loading control (the experiment presented here is representative of at least 3 independent experiments).

### Pharmacological inhibition of LAT1 impairs MB cell proliferation, survival and migration abilities

We next assessed the effect of JPH203-induced LAT1 pharmacological inhibition on cell proliferation and cell viability. Two concentrations of the inhibitor (20 and 30μM) strongly decreased the proliferation of HD-MB03 and DAOY cell lines (Fig.2A). Moreover, while having a cytostatic effect at 20μM, JPH203 was cytotoxic at 30μM in both cell lines (Fig. 2B). This effect was stronger in HD-MB03 (30%) than in DAOY cells (7%) suggesting that the HD-MB03 cell line, belonging to the most aggressive subgroup of MBs and expressing the highest level of LAT1/CD98 complex (Fig. 1A), is also the most sensitive to LAT1 inhibition. The effect of 30μM of JPH203 was tested on murine primary cortical neurons (PCN) and non-tumoral cerebellar astrocytes (C8-D1A). The treatment had no significant effect on PCN viability and only slightly impaired astrocyte viability (Fig. 2C. The effect of JPH203 was then tested on spheroids generated with HD-MB03 and DAOY cells to assess the effect of LAT1 inhibition on the 3-dimensional (3D) growth of tumor cells. As found in 2D, JPH203 completely abolished HD-MB03 and DAOY spheroid growth at two different concentrations (Fig. 2D and E). These results demonstrate that LAT1 activity is crucial for MB cell proliferation and survival.

**Figure 2:**
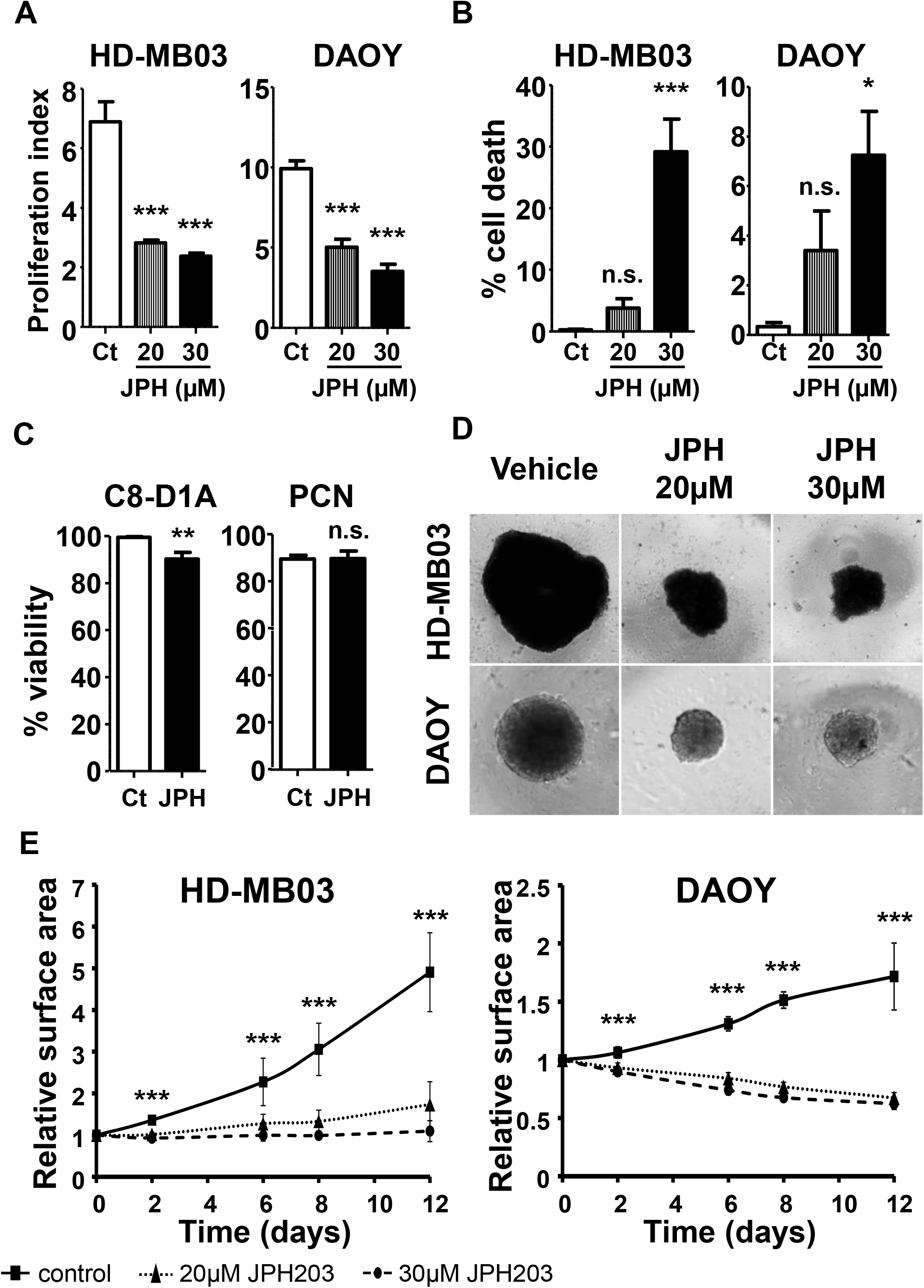
LAT1 inhibition selectively impairs growth and decreases viability of MB cells. (A) Proliferation assay of HD-MB03 and DAOY cells treated or not with 20 or 30μM of JPH203 for 72 hours. Control (Ct) conditions were treated with DMSO for the same time period. (B) Viability assay of HD-MB03 and DAOY cells treated or not with 20 or 30μM of JPH203 for 72 hours. Control (Ct) conditions were treated with DMSO for the same time period. (C) Viability assay of murine primary astrocytes (C8-D1A) and primary cortical neurons (PCN) treated with 30μM of JPH203 or DMSO (control, Ct) for 72 hours. (D) *In vitro* 3-D growth assay of HD-MB03 and DAOY cells. Spheroids generated with HD-MB03 or DAOY cells were treated with 20 or 30μM of JPH203 or DMSO (control, Ct) for 12 days. (E) Spheroid growth measurements over time. The surface area of spheroids was measured at the indicated time points and normalized by the initial size of the control (■), 20μM JPH203 (▲) and 30μM JPH203 (●) spheroids. Data points indicate mean +/- SEM from at least three independent experiments (***: p<0.001; **: p<0.01; *: p<0.05; n.s.: p>0.05, Student’s t-test).

Finally, we tested the effect of LAT1 on the migration capacities of MB cells using a scratch assay on a confluent layer of DAOY cells. JPH203 treatment resulted in a 25% reduction of scratch closure after 8 hours compared to cells treated with DMSO (Fig. 3A and B). This suggests that LAT1 activity is required to sustain MB cell motility. Altogether, these findings demonstrate that LAT1 through its EAA transport activity promotes some of the key malignant features of MB cells while bearing seemingly no or low toxicity towards healthy cerebral cells.

**Figure 3:**
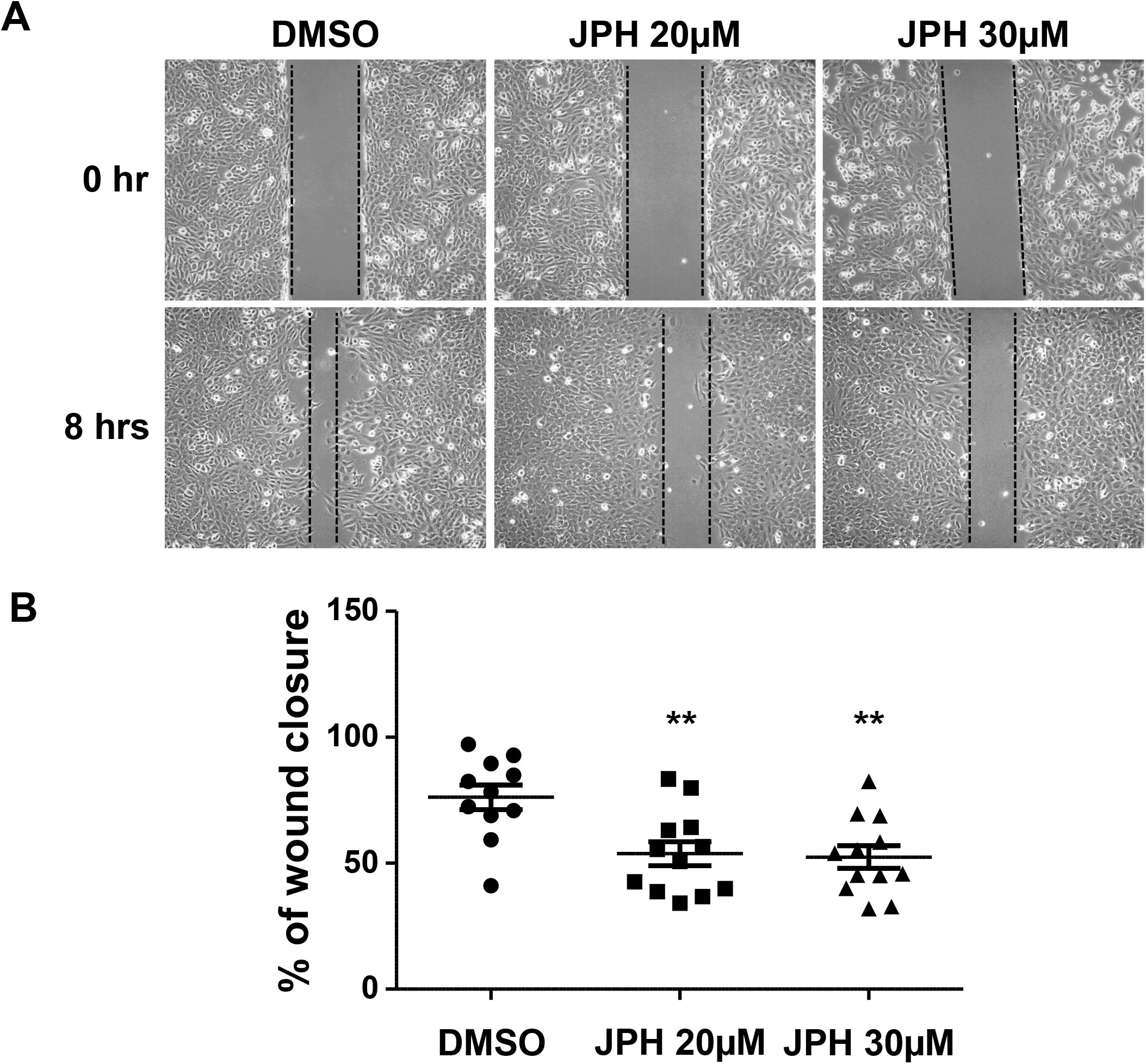
LAT1 inhibition impairs MB cell migration. (A) Scratch-wound assay on a DAOY cell confluent layer treated or not with JPH203 at the indicated concentrations and on which a “mechanical” wound was made at time 0. (B) Dot plot showing quantitative determination of wound closure after 8 hours in the absence or presence of the indicated concentrations of JPH203. The results are expressed as the percentage of closure of each individual wound relative to time 0. The bars represent the mean +/- SEM for each experimental condition (**: p<0.01, Student’s t-test).

### Chronic treatment of MB cells with JPH203 induces cellular adaptation but no resistance

In order to test the development of resistance mechanisms, HD-MB03 and DAOY cell lines were incubated for more than 120 days in JPH203-supplemented medium (20 and 30 μM). The levels of LAT1 and its chaperone CD98 were dramatically increased in response to chronic exposure to JPH203 (Fig. 4A). Consistently, the mRNA levels of LAT1 and LAT3 were also increased (Fig.S1). Nonetheless, JPH203 still displayed strong anti-proliferative and cytotoxic effects on both cell lines (Fig.4B and C). These results strongly suggest that this 4-month adaptation was insufficient to induce resistance to LAT1 inhibition although the chronic treatment with JPH203 led to upregulation of some components of the amino acid transport machinery.

**Figure 4:**
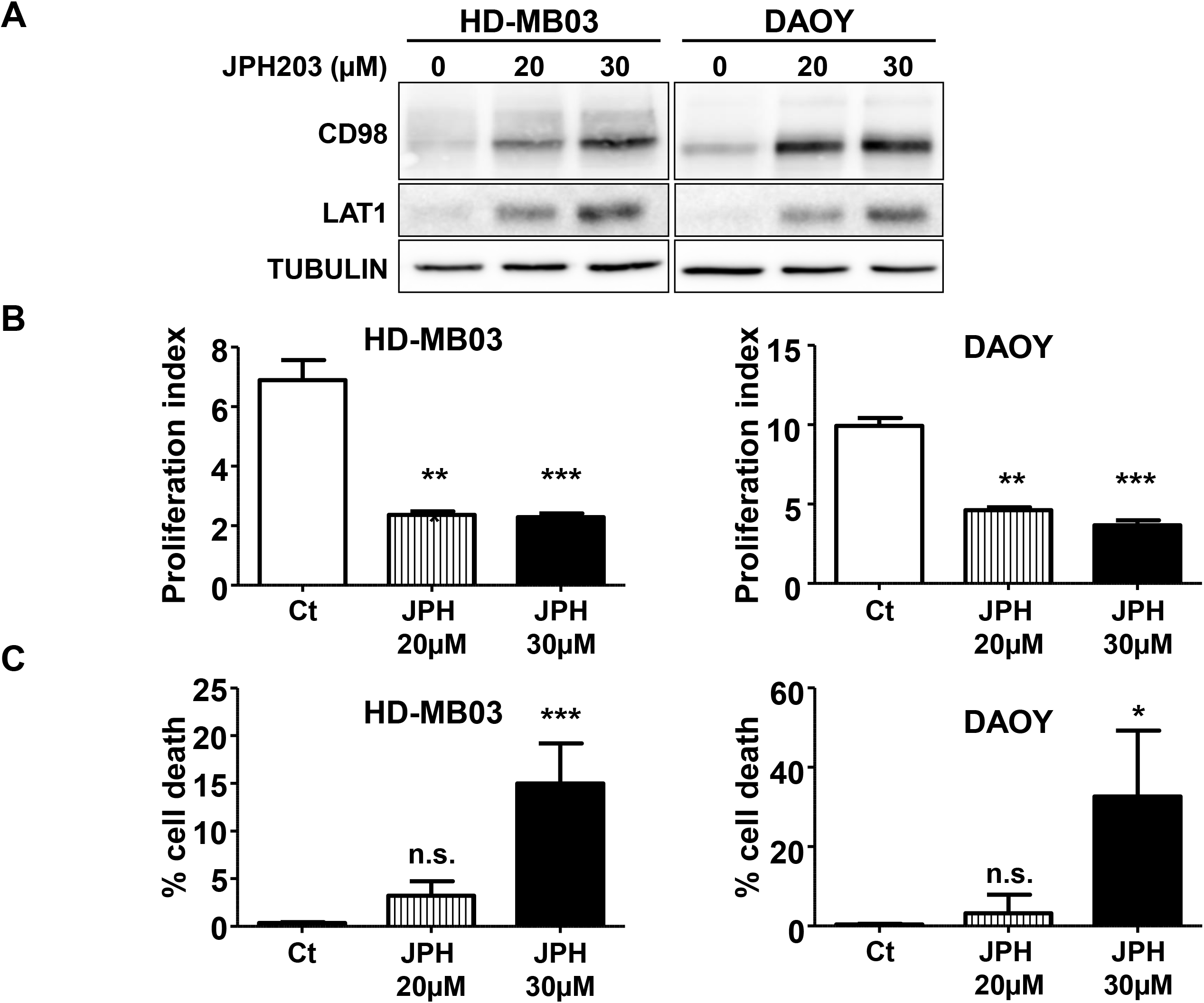
MB cell adaptation to long-term JPH treatment does not cause resistance. (A) The expression levels of CD98 and LAT1 in HD-MB03 and DAOY cells chronically exposed for 120 days to the indicated concentrations of JPH203 were analyzed by immunoblot. Tubulin was used as a loading control. (B) Proliferation assay of HD-MB03 and DAOY cells treated with 20 or 30μM of JPH203 or DMSO (control, Ct) for 120 days. (C) Viability assay of HD-MB03 and DAOY cells treated with 20 or 30μM of JPH203 or DMSO (control, Ct) for 120 days (***: p<0.001; **: p<0.01; *: p<0.05; n.s.: p>0.05, Student’s t-test).

## DISCUSSION

The most aggressive forms of MBs as well as local or metastatic relapses generally show a high degree of resistance to classical treatments. It is therefore crucial to discover new treatments presenting important properties such as selective toxicity towards cancer cells and low toxicity towards normal tissues. JPH203 is a tyrosine analog that inhibits LAT1 with a high specificity as compared to other compounds [22, 26]. The proposed mode of action of this compound is via competition with other substrates of the LAT1 transporters [22]. In the present study, we show that JPH203-induced blocking of LAT1-dependent EAA transport efficiently disrupts AA homeostasis, mTORC1 activity, proliferation and survival of two independent MB cell lines isolated from MBs of different subgroups (high risk Shh group and group 3) [32]. This suggests that JPH203 may represent an efficient treatment for MBs of different genetic backgrounds.

Most current chemotherapies used to treat MB are genotoxic agents [1]. These compounds target all the proliferative cells and are not selective towards cancer cells. This accounts for major side effects observed upon treatment with classical chemotherapeutic agents. The very low toxicity of JPH203 towards primary astrocytes and cortical neurons suggests it may cause little side effects in a clinical setup. The selectivity of JPH203 towards cancer cells can be explained by at least two major mechanisms. First, cancer cells proliferate rapidly as compared to normal cells. They therefore highly rely on amino-acid import as compared to normal cells. This accounts for the strong impairment of cancer cells proliferation when LATs are inhibited. Second, the import of leucine in the cancer cell lines used in the present study as well as other models mostly rely on LAT1 (Fig. 1 B) [23]. Since leucine is an essential amino-acid, these cells cannot grow in the absence of LAT1 activity. Although our results suggest a low toxicity of JPH203 towards normal cells, it is important to mention that the impairment of the transport of branched-chain amino acids at the blood-brain barrier caused by a LAT1 deficiency has also been described to cause autism spectrum disorders in mice [33]. Hence, JPH203 could lead to psycho-cognitive disorders in patients. Yet, according to the results from the first phase 1 clinical trial of JPH203 reported at the American Society of Clinical Oncology (2018 ASCO Annual Meeting), JPH203 is in fact well tolerated in adult patients [34]. Moreover, JPH203 is a non-covalent competitive inhibitor, it should therefore have reversible effects. This may represent an important advantage as compared to the aforementioned genotoxic treatments. Finally, these effects should be weighed against current therapies for MB, generally involving radio- and chemotherapy combinations which are showed to cause major cognitive side effects [1].

The transmembrane transport of JPH203 was demonstrated to be mediated by transporters of the organic-anion-transporting peptides (OATP, SLCO) and the organic anion transporters (OAT, SLC22) families, namely OATP1B1, OATP1B3, OATP2B1 and OAT3 [35]. Interestingly, some of these transporters were also described to be expressed in brain capillary endothelial cells constitutive of the blood-brain barrier (BBR) [36]. This suggests that a route for JPH203 to cross the BBR could be provided by transmembrane transporters, making brain tumors accessible to this compound. Alternatively, delivery of anticancer compounds into the brain can be achieved using Ommaya reservoirs. These medical devices allow intraventricular injection of the drugs, even in very young patients [37].

Development of resistance by cancer cells has proven to be one of the most frequent causes of targeted therapy failure and may account for the low efficacy of therapies targeting the Shh pathway in MB [38,39]. mTORC1 pharmacological inhibition can overcome the acquired resistance to Shh targeted therapy in Shh group MBs [40]. Our study shows that inhibition of LAT1 leads to a strong decrease in mTORC1 activity, suggesting that the use of JPH203 may be relevant to bypass resistance to Shh targeted therapies. We also show that despite an adaptive response via an upregulation of EAA transporter expression, MB cells treated for several months with JPH203 never acquired the capacity to overcome the cytotoxic and cytostatic effects of this compound. This result suggests that MB cells may not have the ability to adapt to EAA import inhibition and subsequent mTORC1 inactivation. Even if encouraging, our results still need *in vivo* validation. In particular, the ability of JPH203 to pass the blood-brain barrier and its toxicity towards developing organisms still require thorough investigation.

## Supporting information

Fulle length WBs for Fig.1A and 4A

## ACKNOWLEDGMENTS

YC and MPB designed and performed the experiments. YC, MPB and VP analyzed the data. YC and VP wrote the manuscript. SS isolated PCN cells. SG contributed to design the experiments and to the manuscript revisions. YC and ET performed the radioactivity experiments. HE and MFW kindly provided the JPH203 but did not conduct any experiment nor participated to the elaboration of experimental procedures. GP and JP contributed reagents, materials and funding. Besides the financial support of CSM, YC and JP were funded by GEMLUC and MPB, VP and GP by Fondation Flavien. All co-authors contributed in improving the draft manuscript. The authors are grateful to Dr. M. Gettings for editing the manuscript and to all the staff members of Fondation Flavien for their constant efforts and strong support.

---

**Figure S1:**
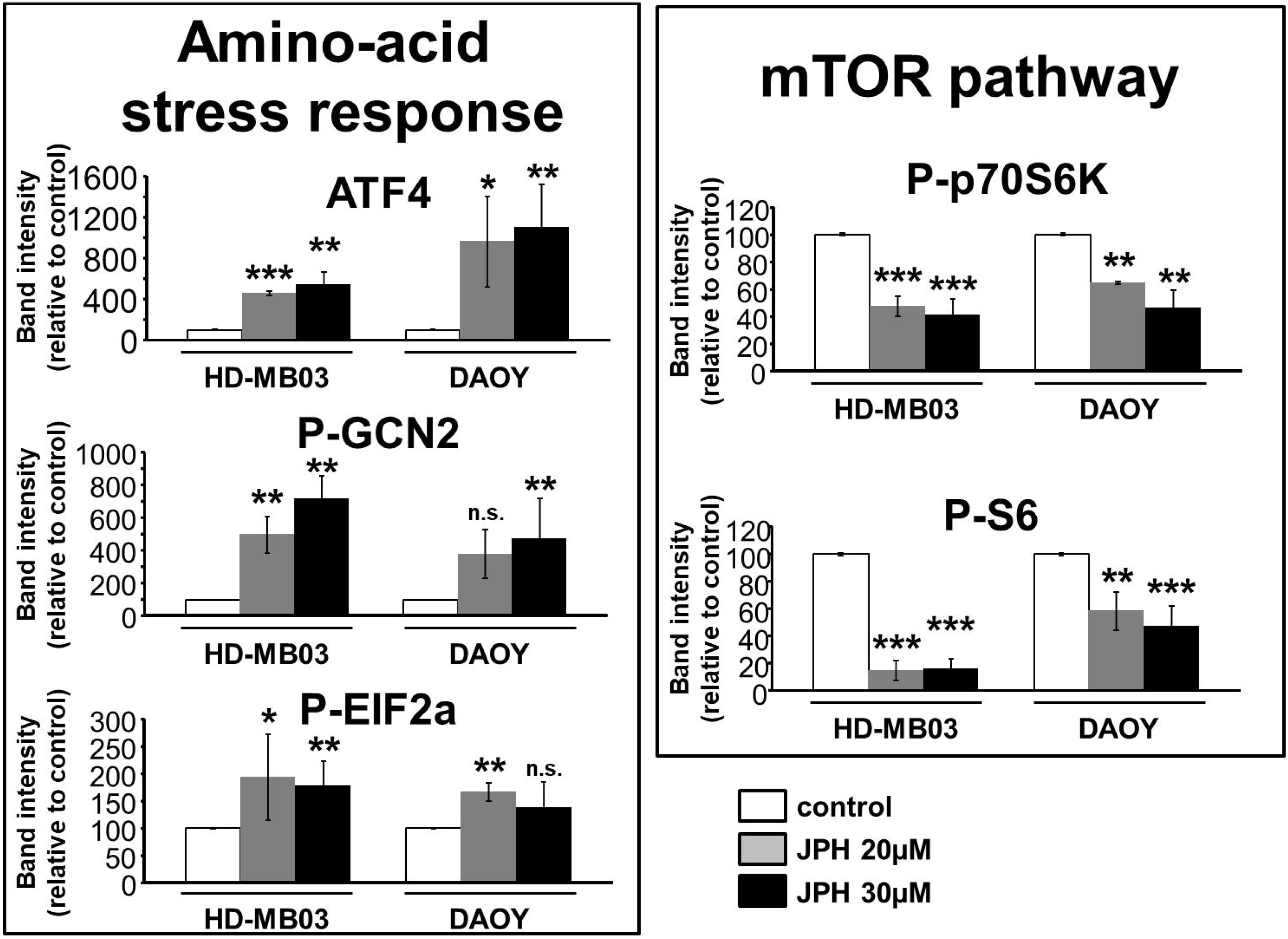
Quantifications of the western-blots for AA sensing pathways. The intensity of the bands corresponding to ATF4 (n=3), phospho-GCN2 (n=4), phospho-EIF2α (n=5), phospho-p70S6K (n=4) and phospho-S6 (n=7) were quantified and normalized to the loading control for ATF4 or to the total protein for the phosphorylated proteins. The results are expressed relative to controls (***: p<0.001, **: p<0.01, *: p<0.1; n.s.: p>0.05; Student’s t-test).

**Figure S2:**
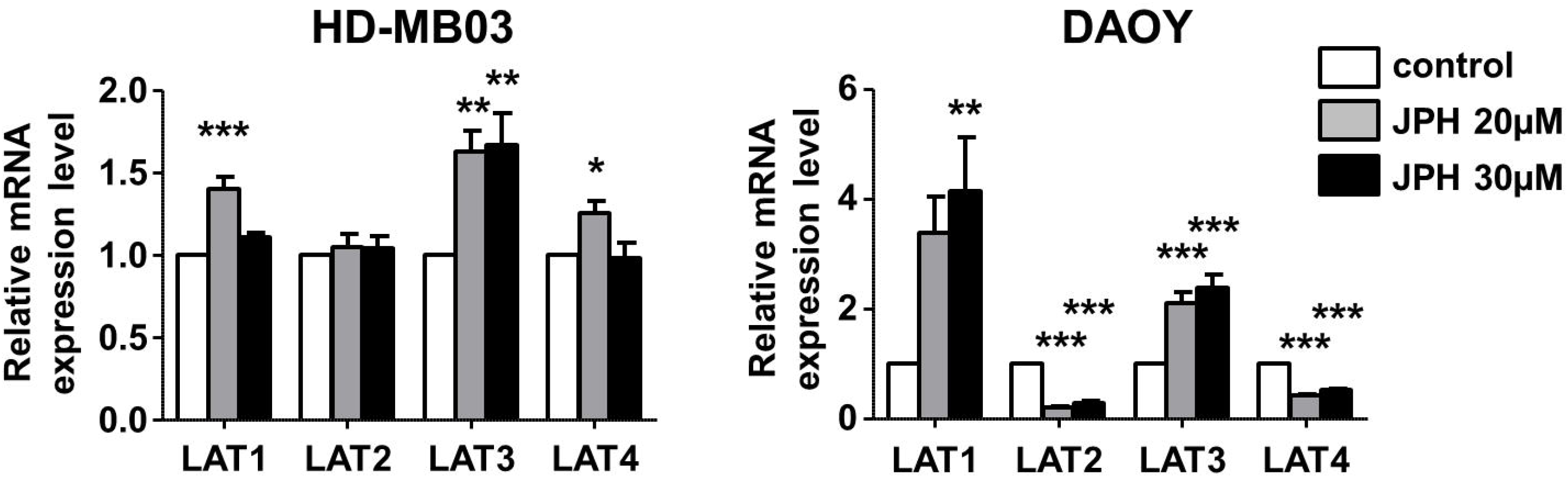
Long-term JPH treatment induces modification in amino acid transporter
gene expression in MB cells. Real-time quantitative PCR analyses of LAT family gene expression levels in HD-MB03 and DAOY cells chronically exposed for 120 days to the indicated concentrations of JPH203. The results are expressed relative to control conditions and were obtained from at least three independent experiments (***: p<0.001, **: p<0.01, *: p<0.1, Student’s t-test).

