## Supplementary figures and images for "Inhibition of the amino-acid transporter LAT1 demonstrates anti-neoplastic activity in medulloblastoma"

### Fulle length WBs for Fig.1A and 4A

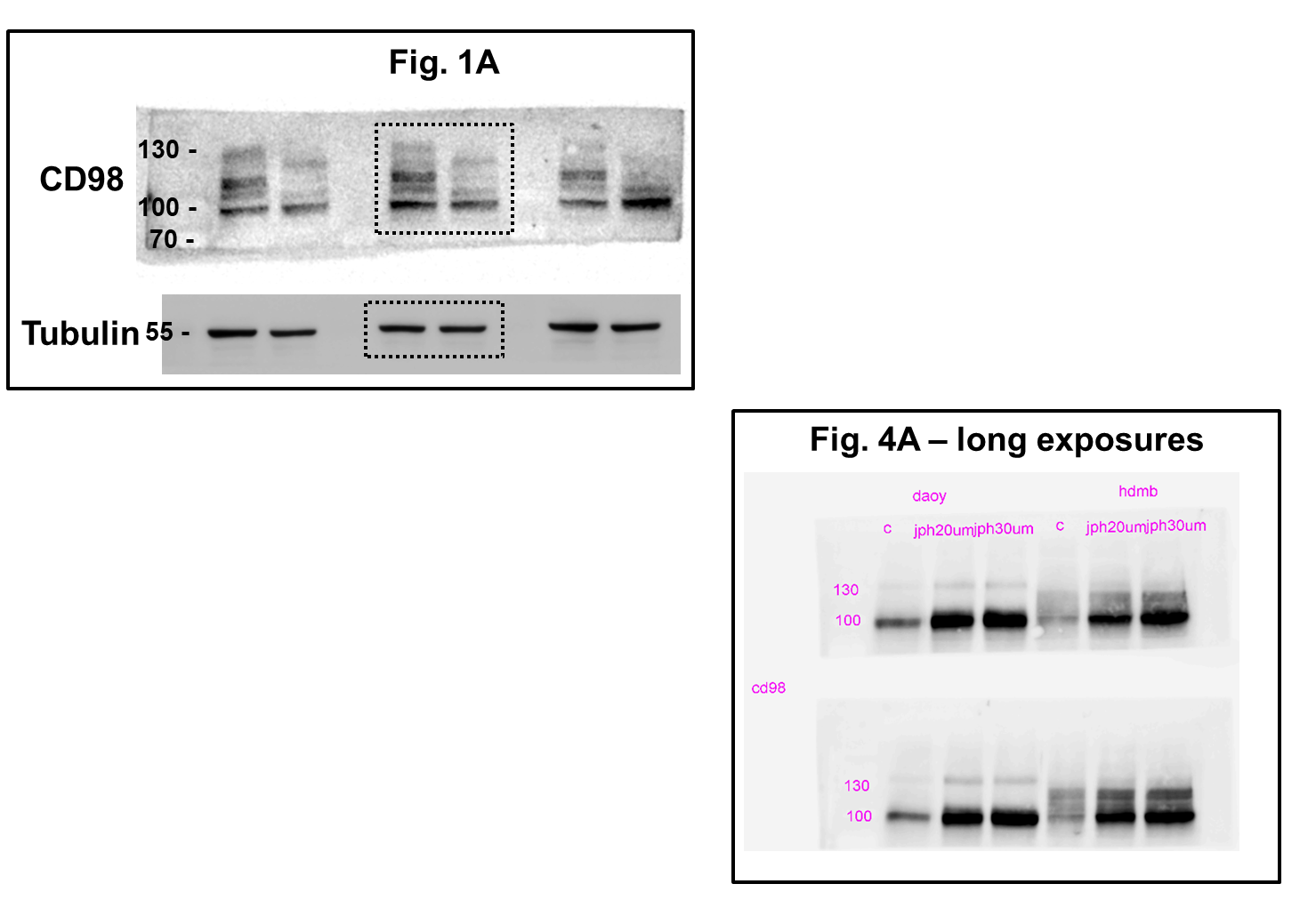
